# BRIX1 Promotes Hepatocellular Carcinoma Progression via the MAPK/ERK Pathway and Serves as a Prognostic Biomarker

**DOI:** 10.64898/2026.08.26.747409

**Authors:** Xianzhu Pan, Xiaoqing Wang, Yuanqin Zhou

## Abstract

Hepatocellular carcinoma (HCC) is particularly aggressive and difficult to treat. Due to the lack of early clinical diagnosis and the unsatisfactory clinical treatment effect, it is particularly important to identify novel markers that can predict tumor behavior in HCC. biogenesis of ribosomes BRX1 (BRIX1) is abundant in various tissues of the human body. However, the regulatory mechanisms and its role in various tissues are not fully understood. Here, we analyzed the expression pattern of BRIX1 in HCC from public gene expression databases and tissue samples from clinical HCC. We confirmed that BRIX1 was upregulated in both HCC cell lines and HCC paraffin section samples. BRIX1 depletion significantly dicreased the capacity of cells to grow and migrate in vitro, and knockdown BRIX1 suppressed tumor growth in xenograft tumor model. Mechanistically, BRIX1 depletion suppressed the MAPK/ERK pathway, as reflected by reduced phosphorylated ERK (p-ERK) levels. In summary, we provide a rational clue for the further investigation of BRIX1 as an invaluable biological marker for diagnosing and predicting prognosis of patients with HCC.

## 1. Introduction

Hepatocellular carcinoma (HCC) accounts for approximately 80–85% of all primary liver malignancies and stands out as the most aggressive and treatment-resistant subtype(1). Although surgical resection combined with systemic therapies has substantially improved outcomes for patients diagnosed at an early stage, HCC continues to pose a major threat to public health, largely due to its high perioperative risks and frequent postoperative recurrence(2–4). The development of HCC typically begins with uncontrolled cellular proliferation, followed by stepwise progression from carcinoma in situ to invasive disease and eventually to distant metastasis. Owing to its rapid progression and the incomplete understanding of its underlying mechanisms, the majority of patients are already at intermediate or advanced stages when first diagnosed, thus missing the window for curative intervention. Moreover, the etiology of HCC is remarkably complex, involving intricate and dynamic interactions among multiple genes at various regulatory levels(5–7). Given the considerable heterogeneity observed across clinical courses, identifying biomarkers that can reliably predict tumor biological behavior is of great clinical value. Such markers are not only useful for diagnosis, disease staging, treatment response evaluation, and detection of recurrence or distant spread, but also contribute to refining prognostic assessment and guiding clinical decision-making. Over the years, a number of candidate biomarkers have been proposed as indicators of HCC progression and aggressiveness; however, most have failed to demonstrate practical clinical utility(8–11). In view of the persistent challenges in early detection and effective prevention of liver cancer, the discovery of robust and actionable biomarkers remains an urgent priority[12]. Furthermore, systematic screening for gene network alterations associated with malignant phenotypes and disease progression may offer a promising avenue to decipher the key molecular events driving HCC pathogenesis, potentially revealing novel therapeutic targets and paving the way for innovative treatment strategies.

BRIX1 (BXDC2) is a conserved nucleolar protein essential for 60S ribosomal subunit assembly(13). Located at 5p13.2, its gene product contains a Brix domain that facilitates RNA binding and is critically involved in rRNA processing and ribosome formation. Recent studies underscore its oncogenic potential: in HCC, BRIX1 upregulation fuels disease progression by promoting ribosome biogenesis and altering alternative splicing through the mTORC1-SP1 axis, and its expression forms part of a prognostic signature linked to poorer outcomes. In colorectal cancer, BRIX1 further drives glycolysis via GLUT1 translation and supports tumor growth by impairing p53 function(14, 15). Nevertheless, the clinicopathological significance of BRIX1 in primary HCC tissues and its independent prognostic utility have yet to be established through direct clinical evaluation.

In the present investigation, we first performed systematic bioinformatic analyses based on the TCGA dataset and subsequently extended our evaluation to an independent clinical cohort of surgically resected HCC specimens, directly examining the correlation between BRIX1 expression and patients’ prognostic outcomes. These clinical observations prompted us to explore the biological functions of BRIX1 in HCC through a series of in vitro functional assays, which confirmed its pro-proliferative and pro-migratory properties. To further substantiate these cellular findings under physiologically relevant conditions, we conducted xenograft tumor models in nude mice, demonstrating that BRIX1 knockdown significantly impeded tumor expansion in vivo. We propose that BRIX1 not only holds promise as a prognostic biomarker for risk stratification but may also represent a potential therapeutic target, offering a rationale for future precision oncology strategies in HCC management.

## 2. Materials and Methods

### 2.1. Data retrieval and preprocessing

Transcriptomic data for HCC cohorts were obtained from the TCGA (TCGA-LIHC) and GEO repositories (https://www.ncbi.nlm.nih.gov/gds/) under accession numbers GSE54236, GSE36376, GSE57957, GSE76427, GSE38174, and GSE25097(16–19). While protein-level expression data were retrieved from the CPTAC database(20). For the TCGA-LIHC and CPTAC datasets, raw read counts were normalized and subsequently transformed using log2 scaling. For the GEO datasets, probe-level signal intensities underwent the same log2 transformation to ensure comparability across platforms. Differential expression analysis between tumor and adjacent non-tumor tissues was performed using Student’s t-test, implemented in SPSS version 20.0. A two-sided P value < 0.05 was considered statistically significant.

### 2.2. Patient tissue sample collection

HCC and paired adjacent non-tumor tissues were collected from patients who underwent surgical resection at the Affiliated Hospital of Anhui Institute of Medicine between 2021 and 2025. Post-resection specimens were fixed, paraffin-embedded, and stored at 4 °C until further use. Relevant clinicopathological parameters, including age, sex, histological grade, depth of invasion, tumor size, lymphatic metastasis, and TNM stage, were extracted from medical records. None of the enrolled patients had received neoadjuvant chemotherapy or radiotherapy prior to surgery. The study protocol was reviewed and approved by the Ethics Committee of Anhui Institute of Medicine. Individual patient data were analyzed anonymously, and the requirement for informed consent was waived by the ethics committee. The study complied with the Declaration of Helsinki.

### 2.3. Survival analysis

Overall survival (OS) was evaluated in relation to BRIX1 protein expression levels and the clinicopathological variables summarized in Table 1. Patients with HCC were stratified into high- and low-BRIX1 expression cohorts using the median expression value as the cut-off. The associations between BRIX1 expression and both OS as well as disease-specific survival (DSS) were further validated using the KM Plotter platform for liver cancer (http://kmplot.com)(21). Survival curves were compared via the log-rank test, with statistical significance defined as P < 0.05.

**Table 1.**
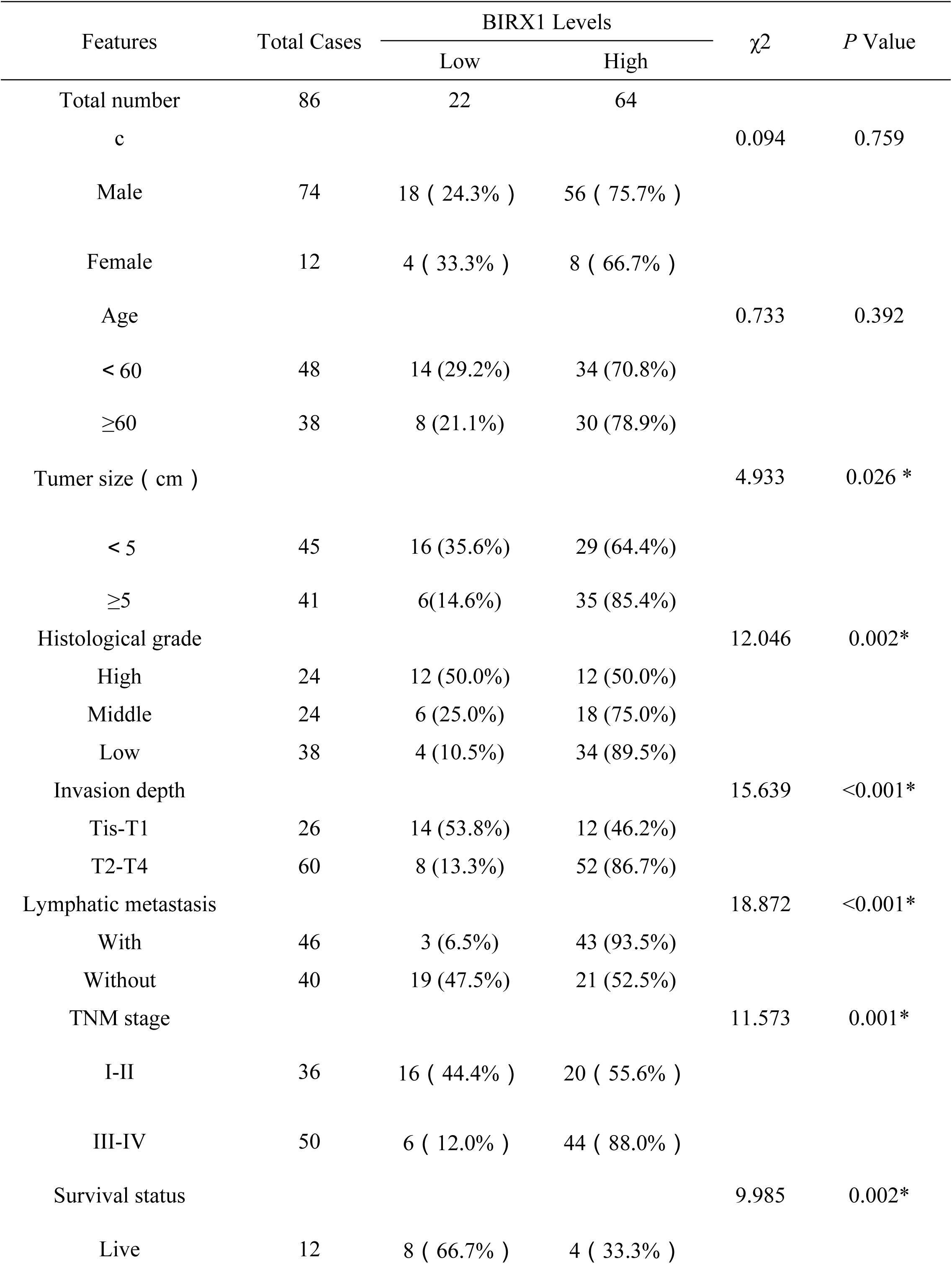
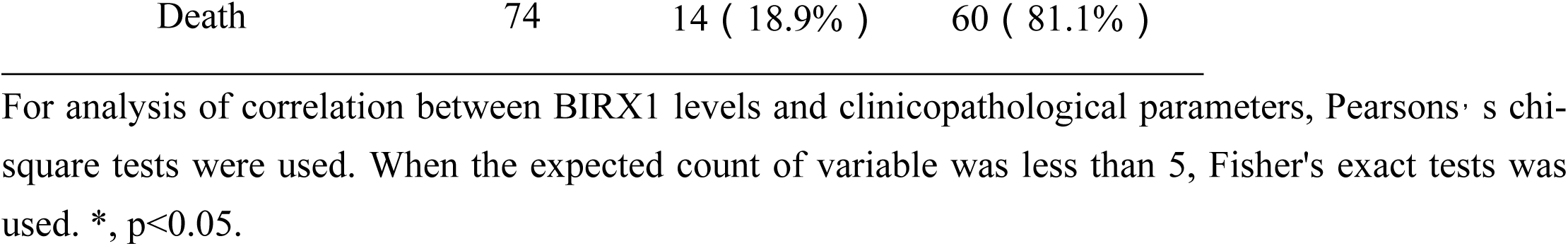
Correlation between BIRX1 protein levels and clinicopathologicl parameter 86 HCC patients.

| Features | Total Cases | BIRX1 Levels | | $\chi^2$ | P Value |
| --- | --- | --- | --- | --- | --- |
|  |  | Low | High |  |  |
| Total number | 86 | 22 | 64 |  |  |
| c |  |  |  | 0.094 | 0.759 |
| Male | 74 | 18 ( 24.3% ) | 56 ( 75.7% ) |  |  |
| Female | 12 | 4 ( 33.3% ) | 8 ( 66.7% ) |  |  |
| Age |  |  |  | 0.733 | 0.392 |
| < 60 | 48 | 14 (29.2%) | 34 (70.8%) |  |  |
| ≥60 | 38 | 8 (21.1%) | 30 (78.9%) |  |  |
| Tumor size ( cm ) |  |  |  | 4.933 | 0.026 * |
| < 5 | 45 | 16 (35.6%) | 29 (64.4%) |  |  |
| ≥5 | 41 | 6(14.6%) | 35 (85.4%) |  |  |
| Histological grade |  |  |  | 12.046 | 0.002* |
| High | 24 | 12 (50.0%) | 12 (50.0%) |  |  |
| Middle | 24 | 6 (25.0%) | 18 (75.0%) |  |  |
| Low | 38 | 4 (10.5%) | 34 (89.5%) |  |  |
| Invasion depth |  |  |  | 15.639 | <0.001* |
| Tis-T1 | 26 | 14 (53.8%) | 12 (46.2%) |  |  |
| T2-T4 | 60 | 8 (13.3%) | 52 (86.7%) |  |  |
| Lymphatic metastasis |  |  |  | 18.872 | <0.001* |
| With | 46 | 3 (6.5%) | 43 (93.5%) |  |  |
| Without | 40 | 19 (47.5%) | 21 (52.5%) |  |  |
| TNM stage |  |  |  | 11.573 | 0.001* |
| I-II | 36 | 16 ( 44.4% ) | 20 ( 55.6% ) |  |  |
| III-IV | 50 | 6 ( 12.0% ) | 44 ( 88.0% ) |  |  |
| Survival status |  |  |  | 9.985 | 0.002* |
| Live | 12 | 8 ( 66.7% ) | 4 ( 33.3% ) |  |  |

For analysis of correlation between BIRX1 levels and clinicopathological parameters, Pearsons' s chi-square tests were used. When the expected count of variable was less than 5, Fisher's exact tests was used. \*, $p < 0.05$ .

### 2.4. Cell culture

Human liver cell lines, including the normal line HL02 and the malignant lines Hep3B, Huh7, HepG2, and PLC, were utilized in this study. All cells were cultured in high-glucose Dulbecco’s modified Eagle’s medium (DMEM; Gibco, Gaithersburg, MD, USA) supplemented with 10% fetal bovine serum (FBS; Lonsera, South America) and 1% penicillin/streptomycin. Cultures were maintained at 37 °C in a humidified atmosphere containing 5% CO₂ within a sterile cell culture incubator (Thermo Fisher Forma 3, USA). Cells in the logarithmic growth phase were harvested for subsequent experiments.

### 2.5. RNA extraction and quantitative real-time PCR (qRT-PCR) detection

Total RNA was extracted from cultured cells using TRIzol reagent (Invitrogen, #15596026) and the TRIzol kit (Invitrogen, #10296010), following the manufacturer’s recommended protocol. For cDNA synthesis, 2 µg of purified RNA was reverse-transcribed using the RT reagent kit (RR037A, Takara, USA) under the following thermal conditions: 42 °C for 30 min, 85 °C for 5 min, and a final hold at 4 °C. Quantitative real-time PCR was subsequently carried out with the SYBR Green qPCR Mix (RR391S, Takara, USA) on a suitable detection system. Gene expression levels of target transcripts were normalized to the internal reference gene GAPDH. Each sample was analyzed in triplicate. Relative mRNA abundance was calculated using the 2⁻ΔΔCT method. The primer sequences employed for qRT-PCR were as follows: BRIX1-F: 5′-TCGGCGTGTCATAAGATCCATCAC-3′, BRIX1-R: 5′-AGTGGGATCATGTGGAAGAAGAGTC-3′, GAPDH-F: 5′-AATGGCAGCAGGCACAAGTACC-3′, and GAPDH-R: 5′-CAAGGGCACAGAGACTAGCGTAATG-3′.

### 2.6. Construction of stable cell lines

Lentiviral vectors carrying short hairpin RNA (shRNA) targeting BRIX1 (shRNA-BRIX1-1 and shRNA-BRIX1-2) or a non-targeting control (shNC) were obtained from Gene Han Bio Company (Shanghai, China). To evaluate the effects of BRIX1 silencing, Hep3B and HepG2 cells were transduced with these lentiviral particles at a multiplicity of infection (MOI) of 20, in the presence of polybrene (2 μg/mL; Han Bio). After 48 hours of co-incubation, successfully transduced cells were selected by continuous exposure to puromycin (2 μg/mL) over a 14-day period. Resistant clones were then manually picked and expanded in fresh medium. Knockdown efficiency was confirmed by qRT-PCR and Western blotting. The shRNA sequences used for BRIX1 targeting were as follows: shRNA-BRIX1-1 sense, 5’AACAGCTTGTGGATTGCATAGGCCA-3’, antisense: 5’-TGGCCTATGCA ATCCACAAGCTGTT-3’; shRNA-BRIX1-2 sense: 5’-TTCTCACTGGCGCAAACAGCTTG TG-3’ and antisense: 5’-CACAAGCTGTTTGCGCCAGTGAGAA-3’.

### 2.7. Cell counting kit-8 (CCK-8) assay

Cell proliferation was assessed using the CCK-8 reagent (MCE, China). Briefly, cells were seeded into 96-well plates at a density of 2,000 cells per well and cultured followed by a 2-hour incubation at 37 °C. At 24-hour intervals over a 72-hour period, CCK-8 solution (10 µL) was added to each well containing 100 µL of DMEM, followed by incubation according to the manufacturer’s instructions. Absorbance was then measured at 450 nm using a microplate reader. Each experimental condition was tested in five technical replicates to ensure reproducibility. Absorbance values obtained at successive time points were plotted to generate cell proliferation curves.

### 2.8. Migration assay

Cell migration was evaluated using Transwell chambers equipped with 8-µm pore-size membranes in 24-well plates (BD Biosciences). Briefly, 4 × 10⁴ cells suspended in 200 µL of serum-free medium (SFM) were seeded into the upper chamber. The lower chamber was filled with complete medium containing 10% fetal bovine serum (FBS) as a chemoattractant. After 18 hours of incubation at 37 °C, non-migrated cells on the upper surface of the membrane were gently removed with a cotton swab. Cells that had migrated to the lower surface were fixed with crystal violet solution, stained, and then counted under a light microscope (BX43, Nikon, Japan ) in five randomly selected fields per well, and quantified using ImageJ software in five randomly selected fields per well.

### 2.9. Western Blot analysis

Total protein was extracted from cultured cells using RIPA lysis buffer (Beyotime, Shanghai, China) supplemented with protease inhibitors. Protein concentrations were determined with a bicinchoninic acid (BCA) assay kit. Equal amounts of protein were separated by SDS-PAGE at 120 V for 90 minutes and subsequently transferred onto polyvinylidene difluoride (PVDF) membranes (Millipore, Boston, MA, USA). Membranes were blocked with 5% bovine serum albumin (BSA) in Tris-buffered saline containing 0.05% Tween-20 (TBST) for 2 hours at room temperature, followed by overnight incubation at 4 °C with primary antibodies. After three washes with TBST, the membranes were incubated with horseradish peroxidase (HRP)-conjugated secondary antibodies (1:8000 dilution; ABclonal, China) for 2 hours at 37 °C. The following primary antibodies were used: anti-BRIX1 (1:1000; ABclonal, A14481), anti-p-ERK1/2 (1:1000; CST, 4377), anti-ERK1/2 (1:1000; CST, 4696), and anti-GAPDH (1:5000; ABclonal, RP00775). Appropriate HRP-conjugated secondary antibodies, including goat-anti-rabbit IgG and goat-anti-mouse IgG (ABclonal), were applied. Protein bands were visualized using enhanced chemiluminescence (ECL) reagents (Thermo Fisher Scientific, USA) and detected with an ECL Plus Western Blotting Detection System.

### 2.10. Immunohistochemistry (IHC) staining

Paraffin-embedded liver cancer tissues were sectioned and processed for immunohistochemical staining. Briefly, sections were deparaffinized with xylene and rehydrated through a graded ethanol series, followed by antigen retrieval in citrate buffer. Endogenous peroxidase activity was blocked, and the sections were then incubated with primary antibodies overnight at 4 °C, including rabbit anti-BRIX1 polyclonal antibody (1:200; ABclonal, A14481) and rabbit anti-Ki-67 polyclonal antibody (1:200; Abcam, ab15580). After thorough washing, the sections were incubated with HRP-conjugated secondary antibodies, and immunoreactivity was visualized using a DAB Detection Kit (Brown, CTS008). Stained sections were examined under a bright-field microscope (BX43, Nikon, Japan). Immunostaining intensity was scored semiquantitatively as follows: 0, negative; 1, weak; 2, moderate; and 3, strong. A final score of ≥ 2 was defined as high expression for subsequent statistical analysis.

### 2.11. Tumor xenograft model

Male BALB/c nude mice (4 weeks old) were obtained from the Nanjing Model Animal Center and housed under specific pathogen-free conditions. All animal experiments were conducted in accordance with institutional guidelines. For tumor implantation, Hep3B cells were harvested by trypsinization, washed three times with phosphate-buffered saline (PBS), and resuspended in serum-free medium. A total of 3 × 10⁶ cells from either BRIX1-knockdown or control groups were subcutaneously injected into the dorsal flanks of nude mice (n = 6 per group) to establish the xenograft model. Tumor dimensions were measured with vernier calipers every four days post-injection. After 22 days, the mice were euthanized by cervical dislocation, and the tumors were excised for further analysis. All animal procedures were reviewed and approved by the Ethics Committee of the Anhui Academy of Medical Sciences.

### 2.12. Statistical analysis

Statistical analyses were carried out using GraphPad Prism 8 and SPSS (version 22.0; SPSS Inc., Chicago, IL, USA). Associations between BRIX1 expression levels and clinicopathological parameters were evaluated using the chi-square test. Survival curves were generated using the Kaplan-Meier method and compared with the log-rank test. All data are presented as mean ± standard deviation (SD) from at least three independent experiments. Comparisons between two groups were performed using Student’s t-test, while comparisons among multiple groups were assessed by one-way analysis of variance (ANOVA). A P value < 0.05 was considered statistically significant, with significance indicated as follows: *P < 0.05, **P < 0.01, ***P < 0.001, and ****P < 0.0001.

## 3. Results

### 3.1. BRIX1 expression patterns and prognostic significance in HCC

We initially observed that BRIX1 expression was selectively elevated in tumor tissues relative to normal tissues (Supplementary Figure S1). To further characterize its potential role in hepatocarcinogenesis, we interrogated publicly available transcriptomic datasets for BRIX1 expression alterations. Analysis of the TCGA-LIHC cohort and six independent GEO datasets (GSE54236, GSE36376, GSE57957, GSE76427, GSE38174, and GSE25097) revealed significantly higher BRIX1 mRNA levels in HCC tissues compared with adjacent non-tumor liver tissues (P < 0.05; Figures 1A and 1C). Consistently, BRIX1 protein expression was also markedly elevated in CPTAC HCC samples relative to normal controls (Figure 1B). We next assessed the prognostic relevance of BRIX1 using the TCGA-LIHC dataset. Kaplan-Meier survival analysis demonstrated that patients with high BRIX1 expression experienced significantly poorer overall survival (OS; Figure 1D) and disease-specific survival (DSS; Figure 1E) than those with low BRIX1 levels. Collectively, these findings indicate that elevated BRIX1 expression is closely associated with an unfavorable prognosis in HCC patients.

**Figure 1.**
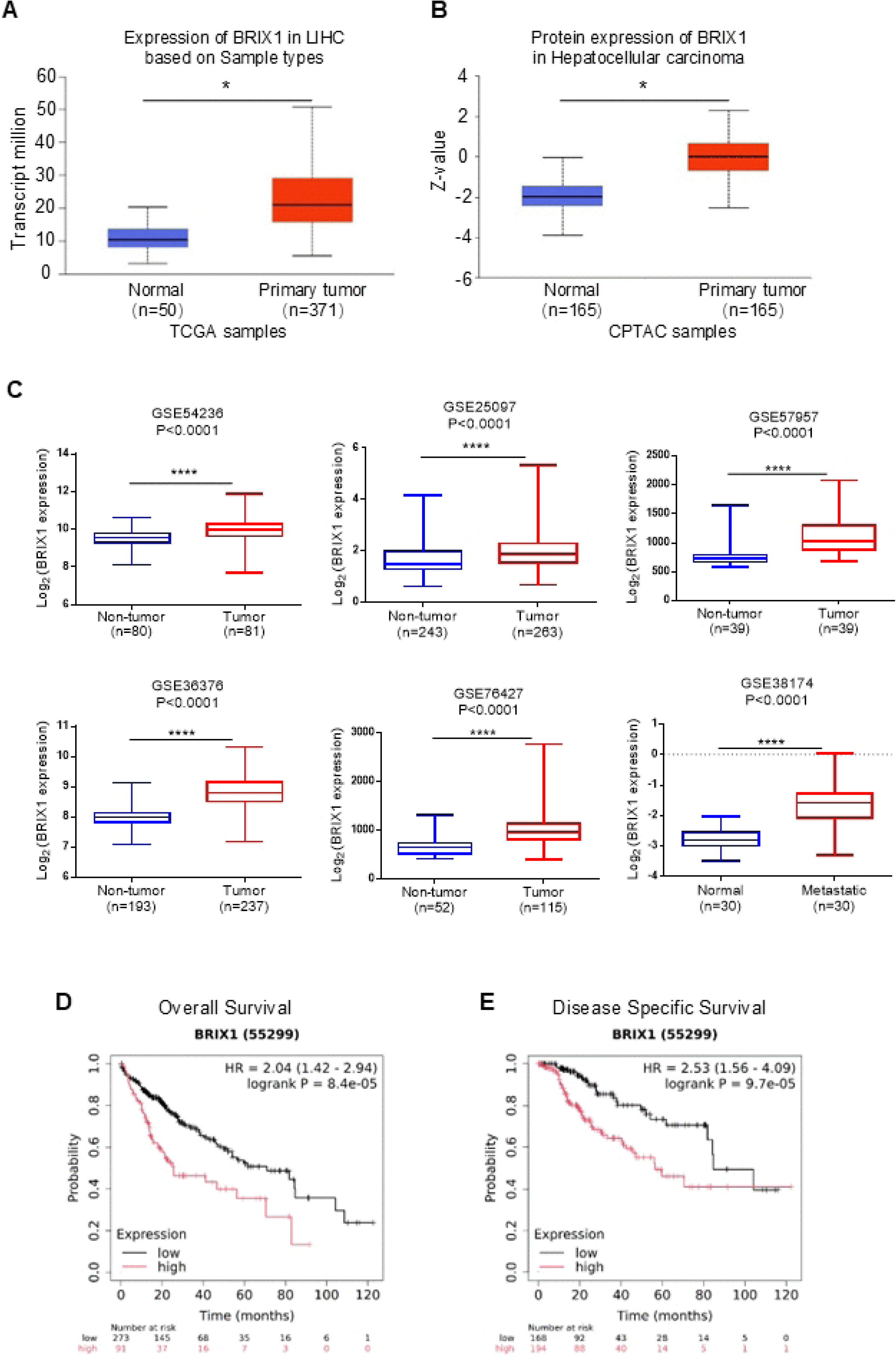
BRIX1 is highly expressed in HCC and correlates with poor patient prognosis. (A) Comparison of BRIX1 mRNA expression levels between HCC tissues and normal liver tissues from the TCGA dataset. (B) BRIX1 protein expression in normal and HCC specimens from the CPTAC dataset. (C) BRIX1 mRNA levels in normal and HCC samples derived from GEO cohorts. (D, E) Kaplan-Meier survival curves showing overall survival (D) and disease-specific survival (E) stratified by BRIX1 expression status. *P < 0.05; ***P < 0.001.

### 3.2. BRIX1 expression is elevated in clinical HCC specimens

To validate the upregulation of BRIX1 observed in public databases, we performed IHC staining on 86 pairs of paraffin-embedded HCC and matched adjacent non-tumor tissues (Table 1). Consistent with the bioinformatics findings, BRIX1 protein levels were markedly higher in tumor tissues than in their adjacent normal counterparts (P < 0.0001, Wilcoxon matched-pairs signed-rank test; Figure 2A). Low BRIX1 expression was observed in 74.4% (64/86) of HCC samples, compared with only 18.6% (16/86) of adjacent normal tissues (P < 0.001; Figures 2B and 2C). IHC analysis further revealed that BRIX1 was predominantly localized to the nucleus in HCC cells (Figure 2C). We next examined BRIX1 expression at both the protein and mRNA levels across a panel of hepatoma cell lines (HepG2, Huh7, Hep3B and PLC) and the normal immortalized hepatocyte line HL02, using western blotting and qRT-PCR, respectively. In line with the tissue data, both BRIX1 mRNA and protein levels were significantly elevated in all hepatoma cell lines relative to HL02 cells (Figures 2E and 2F).

**Figure 2.**
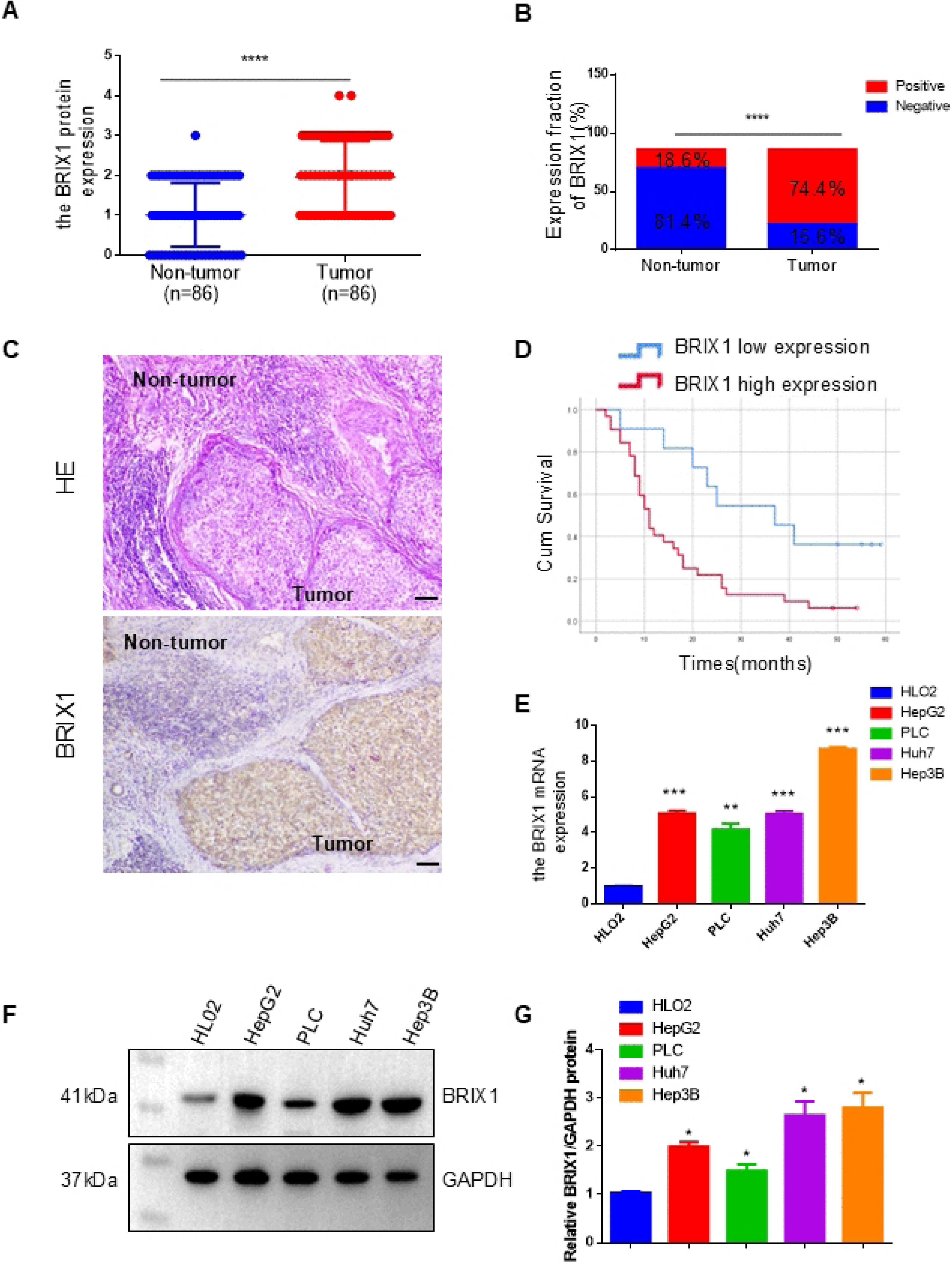
BRIX1 is frequently upregulated in HCC. (A) BRIX1 protein levels in 86 pairs of HCC and matched adjacent non-tumor tissues determined by IHC staining. (B) Quantitative comparison of BRIX1 IHC scores between HCC and adjacent non-tumor tissues (n = 86). (C) Representative IHC images showing BRIX1 expression in HCC tumor and paired adjacent non-tumor samples (scale bar = 200 μm). (D) Kaplan-Meier survival curve depicting overall survival of 86 HCC patients stratified by BRIX1 expression levels; P values were calculated using the log-rank test. (E) BRIX1 mRNA levels in HCC cell lines and the normal immortalized hepatocyte line HL02 measured by qRT-PCR. (F) BRIX1 protein expression in HCC cell lines and HL02 cells assessed by western blotting. *P < 0.05; **P < 0.01; ***P < 0.001.

### 3.3. High BRIX1 expression correlates with tumor progression and poor prognosis in HCC

We next investigated the relationship between BRIX1 expression and clinicopathological features in HCC patients (Table 1). Chi-square test analysis revealed that BRIX1 levels were significantly associated with histological grade (P = 0.002), lymphatic metastasis (P = 0.031), and survival status (P < 0.001; Table 1). Kaplan-Meier survival analysis further demonstrated that HCC patients with high BRIX1 expression had significantly shorter overall survival than those with low BRIX1 expression (P < 0.001; Figure 2D). Moreover, multivariate Cox regression analysis identified BRIX1 expression, lymph node metastasis, tumor size, depth of invasion, and invasion status as independent prognostic factors for overall survival in HCC patients (Table 2).

**Table 2.** Univariate and multivariate analysis of overall survival in 86 patients with HCC by COX regression analysis.

| Variables | Univariate analysis |  | Multivariate analysis |  |
| --- | --- | --- | --- | --- |
|  | HR(95%CI) | *p* value | HR(95%CI) | *p* value |
| Gender | 1.308 (0.671-2.553) | 0.431 | - | - |
| Age | 1.437 (0.932-2.328) | 0.097 | - | - |
| Tumor size (cm) | 2.727 (1.709-4.352) | <0.001* | 2.515 (1.491-4.241) | 0.001* |
| Histological grade (Middle) | 2.646 (1.352-5.178) | 0.005* | 2.225 (1.049-4.719) | 0.037* |
| (Low) | 3.774 (2.031-7.013) | <0.001* | 1.875 (0.872-4.034) | 0.108 |
| Invasion depth | 3.748 (2.139-6.564) | <0.001* | 2.275 (1.067-4.853) | 0.033* |
| Lymphatic metastasis | 2.017 (1.267-3.212) | 0.003* | 0.960 (0.526-1.753) | 0.894 |
| TNM stage | 2.188 (1.357-3.529) | 0.001* | 1.145 (0.648-2.024) | 0.640 |
| BIRX1 expression | 2.976 (1.646-5.380) | <0.001* | 2.155 (1.154-4.024) | 0.016* |

Abbreviations: HR, hazard ratio; CI, confidence interval; \*, $p < 0.05$ .

### 3.4. BRIX1 knockdown suppresses HCC cell proliferation and migration in vitro

Given the elevated BRIX1 expression observed in HCC tissues, we speculated that BRIX1 might contribute to HCC cell growth. To investigate its functional role, we knocked down endogenous BRIX1 in Hep3B and HepG2 cells using lentiviral shRNA vectors. Transfection efficiency was confirmed by qRT-PCR and western blotting, which showed that BRIX1 expression was reduced by approximately 70% in sh-BRIX1-treated cells relative to controls (Supplementary Figures S2a and S2b). CCK-8 assays revealed that BRIX1 knockdown significantly impaired the proliferative capacity of both Hep3B and HepG2 cells (Figure 3A). Given that BRIX1 expression was associated with tumor metastasis in clinical HCC samples, and that metastatic spread accounts for the majority of cancer-related deaths [14], we next assessed the impact of BRIX1 on cell migration. Transwell migration assays demonstrated that BRIX1 depletion markedly reduced the migratory ability of both Hep3B and HepG2 cells compared with controls (Figures 3B and 3C). The MAPK signaling cascade is known to regulate multiple cellular processes, including metabolism, proliferation, survival, metastasis, and angiogenesis in response to extracellular stimuli [15,16]. To determine whether BRIX1 participates in these oncogenic processes, we examined the effect of BRIX1 knockdown on MAPK pathway activity. As shown in Figures 3D and 3E, western blotting revealed that BRIX1 depletion substantially attenuated the phosphorylation level of ERK1/2. Collectively, these findings indicate that BRIX1 promotes HCC progression, at least in part, through activation of the ERK/MAPK signaling axis.

**Figure 3.**
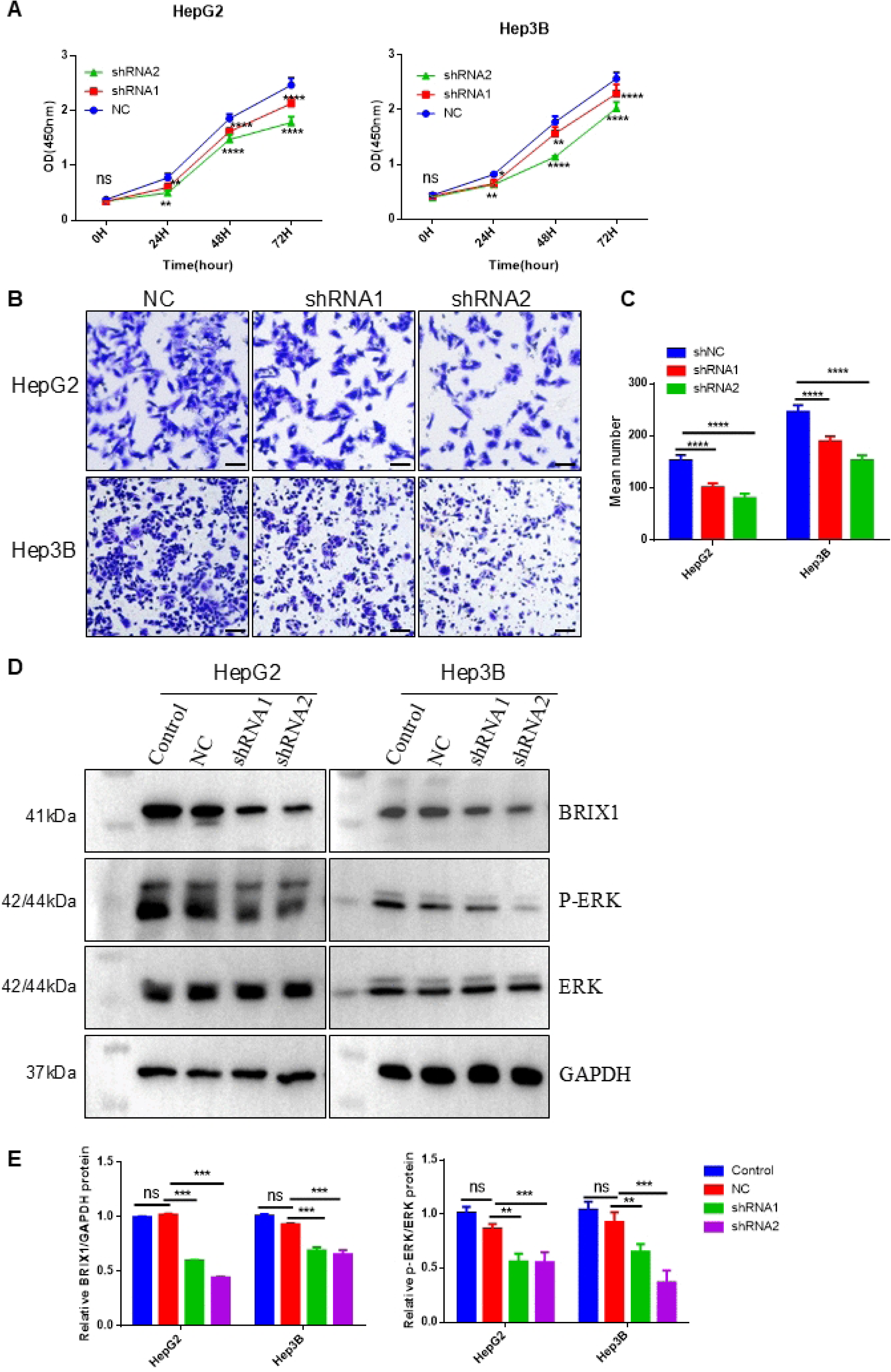
BRIX1 knockdown suppresses HCC cell proliferation and migration in vitro. (A) CCK-8 assays assessing the viability of BRIX1-knockdown Hep3B and HepG2 cells over a 4-day period, with measurements taken daily. (B) Transwell migration assays performed in Hep3B and HepG2 cells transduced with BRIX1-targeting shRNA lentivirus (scale bar = 200 μm). (C) Quantification of migrated cells per field for the two groups. (D) Whole-cell lysates from Hep3B and HepG2 cells transduced with BRIX1 shRNA lentivirus were subjected to immunoblotting with the indicated antibodies; representative blots are shown (n = 3). *P < 0.05; **P < 0.01; ***P < 0.001.

### 3.5. BRIX1 knockdown suppresses tumor growth in vivo

To further validate the tumor-promoting role of BRIX1, we established a xenograft mouse model by subcutaneously injecting Hep3B cells stably transduced with BRIX1-targeting shRNA (sh-BRIX1) or empty lentiviral vector (vector control) into the dorsal flanks of nude mice. Knockdown efficiency was confirmed prior to injection (Supplementary Figure S1B). Following 22 days of tumor growth, BRIX1 knockdown significantly suppressed tumor progression compared with the control group (Figures 4A and 4B). Tumors in the sh-BRIX1 group exhibited markedly smaller volumes (Figure 4B) and lower weights (Figure 4C) than those in the control group. The mean tumor weight was 0.802 g in the knockdown group versus 1.177 g in the control group, corresponding to an inhibition rate of approximately 31.9% (Figure 4C). No significant differences in total body weight were observed between the two groups (Figure 4D), indicating that the observed reduction in tumor growth was not attributable to systemic cachexia. Collectively, these findings demonstrate that BRIX1 plays an essential role in promoting HCC tumorigenesis in vivo.

**Figure 4.**
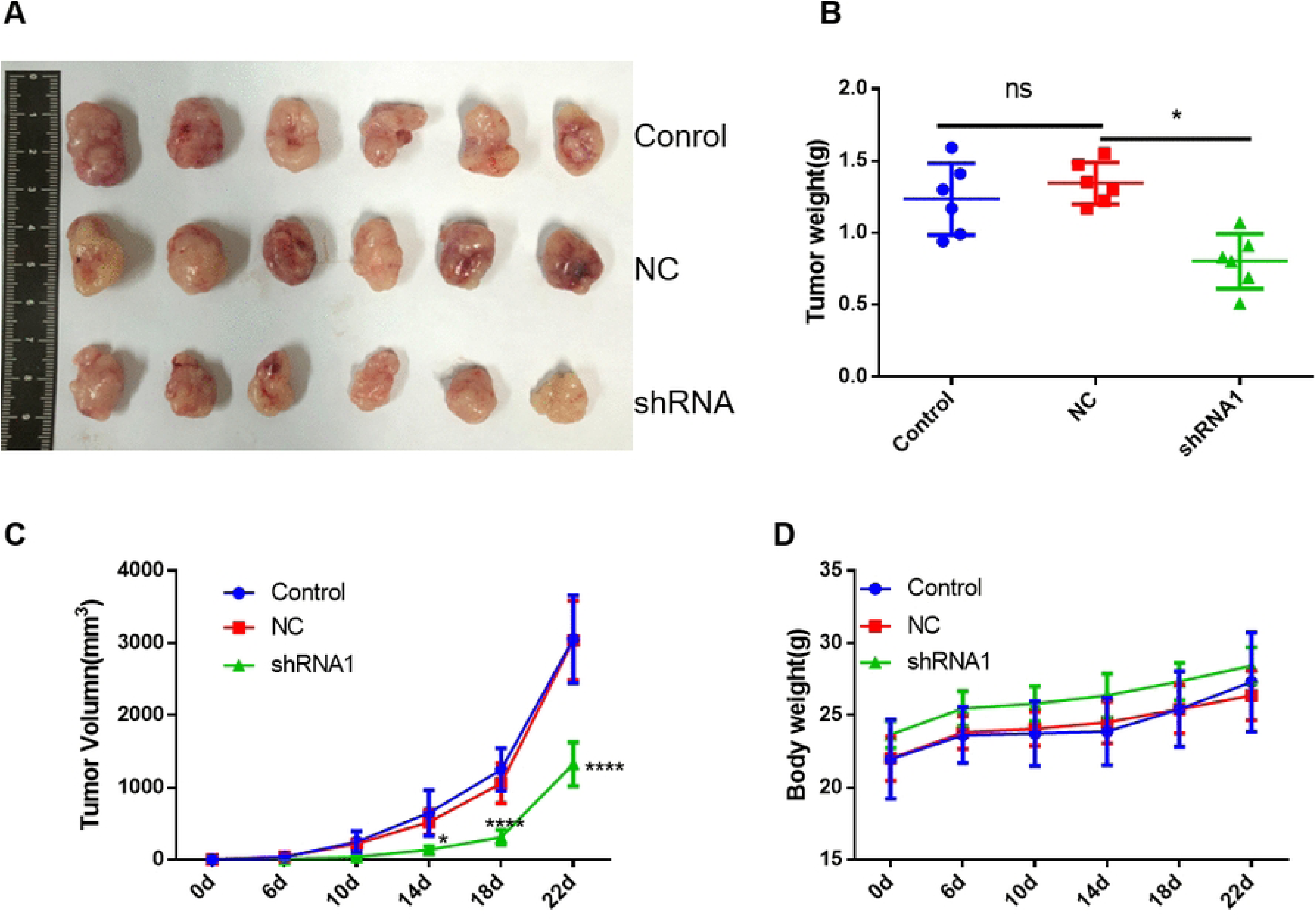
BRIX1 knockdown suppresses tumor growth in vivo. (A) Representative images of xenograft tumors derived from nude mice subcutaneously injected with Hep3B cells, Hep3B cells transduced with BRIX1-targeting shRNA or control vector (3 × 10⁶ cells per injection site). (B) Tumor growth curves showing periodic measurements of tumor volume over time. (C) Tumor weights at the endpoint for each group. (D) Body weight curves of mice across the three groups. *P < 0.05; **P < 0.01; ***P < 0.001.

## 4. Discussion

BRIX1 participates in multiple facets of ribosomal biogenesis and cellular homeostasis(13, 14, 22). Disruption of its normal function may therefore compromise translational fidelity and stress adaptation, ultimately contributing to the aberrant proliferative advantage observed in malignant cells(23). Previous transcriptomic analyses have implicated BRIX1 in the mTORC1-SP1 signaling cascade and alternative splicing regulation(13), and its elevated expression has been associated with unfavorable prognosis in hepatocellular carcinoma patients. Nevertheless, the precise functional repertoire of BRIX1 in cancer—particularly its direct impact on tumor initiation and progression—remains incompletely defined. In this study, we addressed this gap by integrating clinical correlation, in vitro functional assays, and in vivo xenograft models, providing direct evidence that BRIX1 drives HCC aggressiveness and serves as a potential prognostic indicator.

HCC is a complex and multifactorial malignancy, with its pathogenesis closely linked to chronic hepatitis B(24) or hepatitis C virus infection(25), dietary aflatoxin exposure(26), non-alcoholic fatty liver disease(27), diabetes mellitus(28), tobacco smoking(29), and diverse genetic aberrations(30). Despite advances in treatment, the overall prognosis for HCC patients remains discouragingly poor, highlighting an urgent need for more effective therapeutic options(31). In this study, by integrating bioinformatics mining, clinical tissue validation, in vitro functional assays, and in vivo xenograft experiments, we show for the first time that BRIX1 is markedly overexpressed in both HCC tissues and cell lines. This elevated expression was closely associated with adverse clinicopathological features and reduced overall survival, aligning well with findings from public genomic repositories. These observations collectively point to BRIX1 upregulation as a hallmark of HCC and a candidate prognostic indicator. Of note, our results are supported by a recent report showing that BRIX1 is also upregulated in colorectal cancer, where it promotes glycolysis and tumor proliferation(14). Taken together, our work provides compelling evidence that BRIX1 functions as a bona fide oncogenic driver and may represent a viable therapeutic target for HCC intervention.

The oncogenic relevance of BRIX1 appears to extend beyond HCC. In colorectal cancer, for example, BRIX1 overexpression has been reported to enhance glycolytic metabolism and drive tumor growth via selective translation of GLUT1[14], suggesting that its pro-tumorigenic functions may be operative across multiple tumor types. To explore whether BRIX1 exerts a comparable driving effect in HCC, we conducted loss-of-function experiments in Hep3B and HepG2 cells. Our results showed that BRIX1 knockdown markedly reduced cell viability and migratory capacity. These in vitro observations were further validated in vivo using a xenograft mouse model, where BRIX1 knockdown significantly impeded subcutaneous tumor growth. Taken together, these complementary findings from cellular and animal models establish a critical role for BRIX1 in driving HCC progression and metastasis, further supporting its classification as a bona fide oncogenic factor in this disease.

The oncogenic relevance of BRIX1 is not confined to hepatocellular carcinoma. In colorectal cancer, for instance, elevated BRIX1 expression has been shown to enhance glycolytic metabolism and promote tumor growth through selective translation of GLUT1(14), indicating that its pro-tumorigenic functions may extend across diverse cancer types. To determine whether BRIX1 exerts a similar driving role in HCC, we performed loss-of-function experiments in Hep3B and HepG2 cell lines. Our data revealed that BRIX1 knockdownded significantly suppressed cell viability and migration, These in vitro observations were further corroborated by xenograft tumor models in nude mice, wherein BRIX1 knockdown substantially accelerated subcutaneous tumor growth, while its overexpression effectively restrained tumor expansion in vivo. Collectively, these convergent findings from cellular and animal models establish a critical role for BRIX1 in driving HCC progression and metastasis, reinforcing its identity as a bona fide oncogenic factor in this malignancy.

The MAPK/ERK signaling cascade serves as a pivotal nexus integrating extracellular mitogenic stimuli with intracellular transcriptional programs that govern cell proliferation, survival, and motility (32, 33). As a core component of this pathway, ERK1/2 relays signals from receptor tyrosine kinases to downstream effectors, and its sustained hyperactivation has been widely implicated in hepatocarcinogenesis, correlating with aggressive tumor behavior and poor clinical outcomes(34–36). To determine whether BRIX1 exerts its oncogenic effects through this signaling axis, we examined the phosphorylation status of ERK1/2 following BRIX1 manipulation in HCC cells. Our results demonstrated that BRIX1 knockdown markedly reduced the phosphorylated levels of ERK1/2. These findings indicate that BRIX1 functions as a positive upstream regulator of the ERK signaling pathway in HCC cells. Given the well-established role of ERK hyperactivation in driving malignant progression— including enhanced proliferative capacity, resistance to apoptosis, and metastatic dissemination(37–39). our observations suggest that the pro-tumorigenic effects of BRIX1 are, at least in part, mediated through ERK pathway activation. Collectively, these data unveil a previously unrecognized BRIX1-ERK regulatory axis, providing a mechanistic framework for understanding how BRIX1 promotes HCC aggressiveness and highlighting its potential as a therapeutic target for disrupting ERK-driven oncogenic signaling in this malignancy.

Our findings establish BRIX1 as a key oncogenic driver in HCC, yet certain limitations should be acknowledged. Mechanistically, we primarily focused on the ERK pathway, while the upstream regulators of BRIX1 and additional downstream effectors remain to be characterized. Functionally, although we confirmed the effects of BRIX1 on proliferation, migration, and tumor growth, the complete signaling network warrants further dissection. Clinically, our prognostic analyses were based on a single cohort, and multi-center validation would enhance their generalizability. Additionally, the differential response of normal versus malignant cells to BRIX1 perturbation has not been examined. Addressing these issues will be critical for translating our findings toward clinical application.

## 5. Conclusions

In conclusion, our integrative study demonstrates that BRIX1 is significantly overexpressed in HCC, where its elevated expression correlates with aggressive clinicopathological features and poor prognosis, supporting its utility as a prognostic biomarker. Functional assays confirm that BRIX1 promotes HCC cell proliferation, migration, and tumor growth in vivo, while mechanistic dissection reveals that these effects are mediated, at least in part, through positive regulation of the ERK signaling pathway. Collectively, these findings establish BRIX1 as a bona fide oncogenic driver and a potential therapeutic target in HCC. Further investigations into its upstream regulators and multi-center clinical validation will be essential to translate these observations toward clinical application

## Supplementary Materials

**Figure S1.**
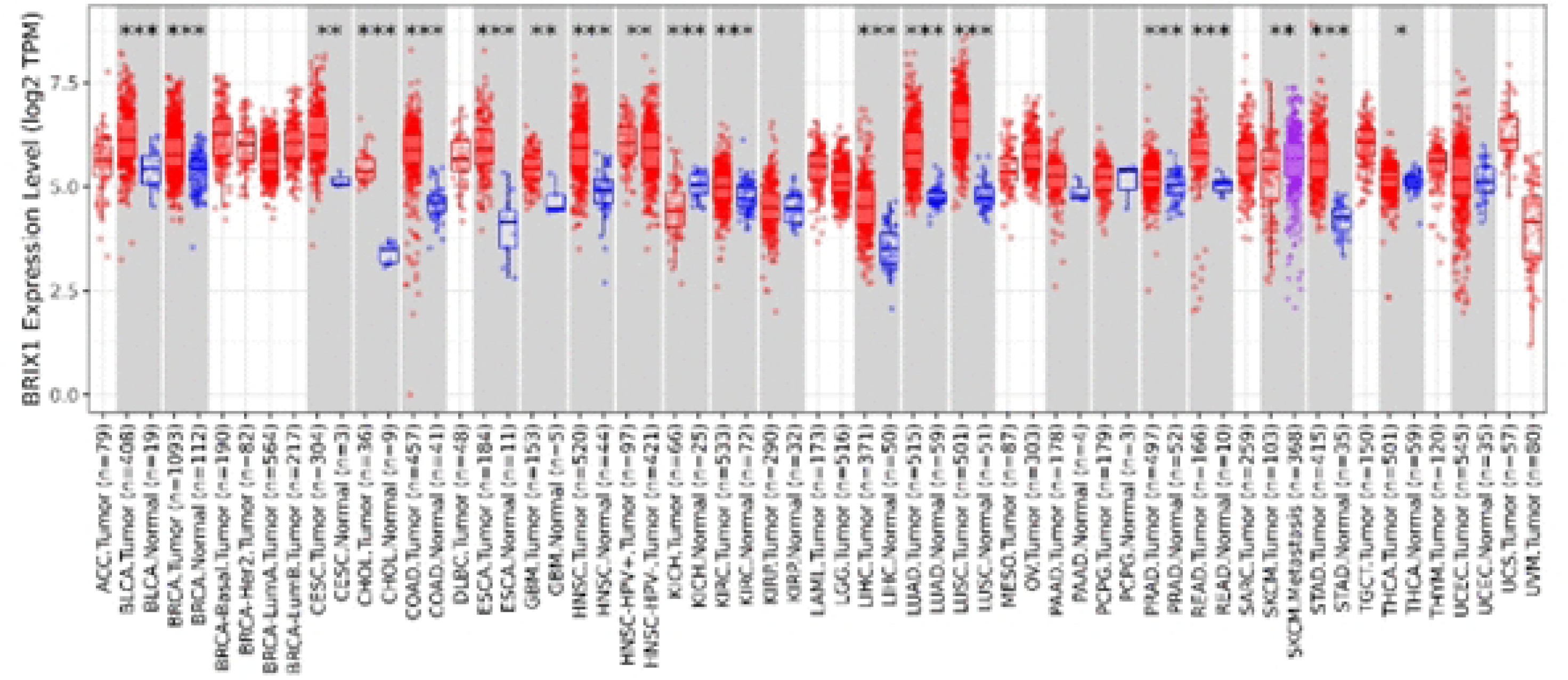
mRNA expression levels of BRIX1 in normal and paired tumor tissues from the TCGA dataset.

**Figure S2:**
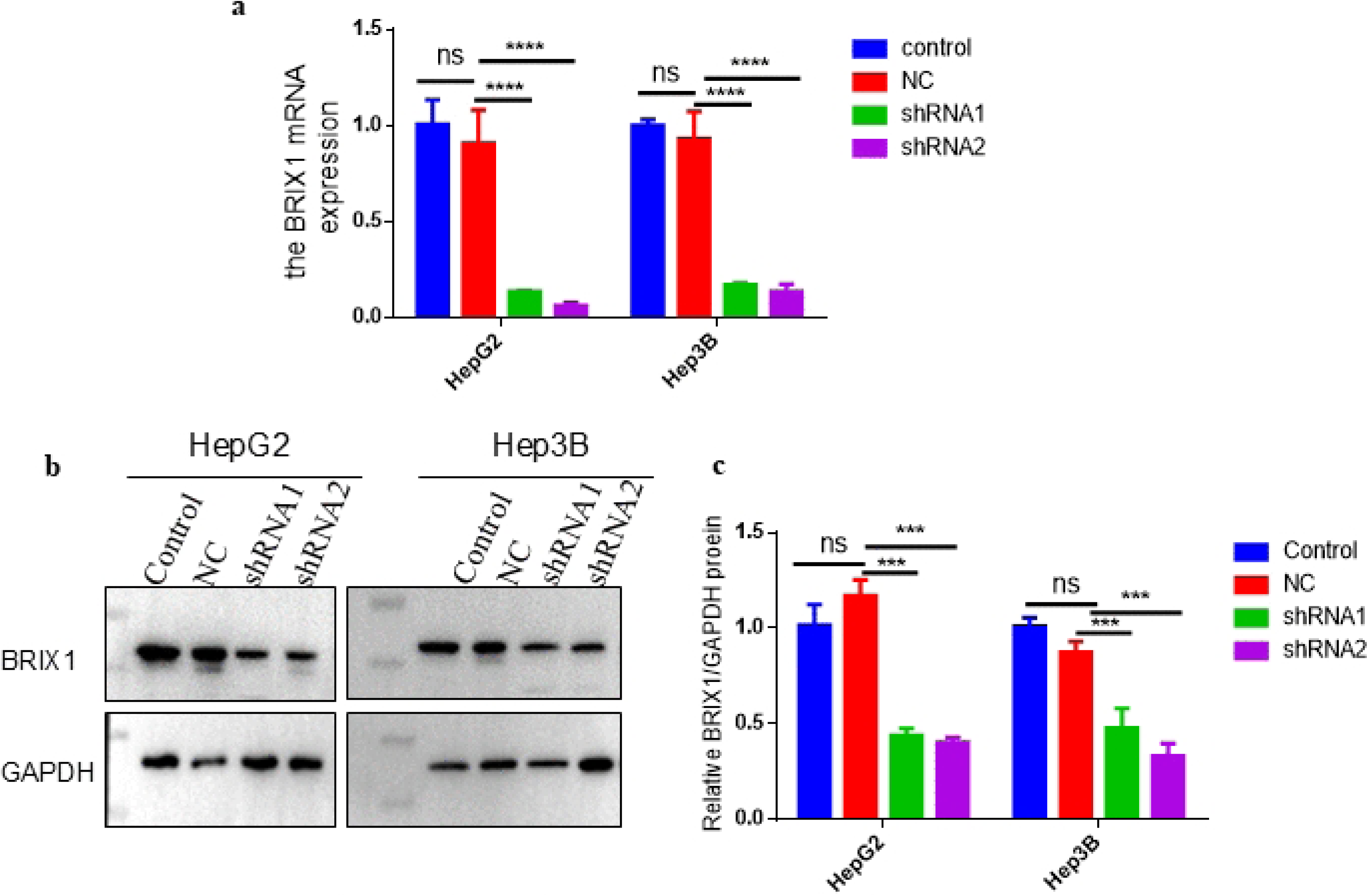
Validation of BRIX1 knockdown efficiency in Hep3B and HepG2 cells. (a) BRIX1 mRNA levels were measured by qRT-PCR following lentiviral transduction with BRIX1-targeting shRNA. (b,c) BRIX1 protein levels were assessed by western blotting under the same conditions. (*Supplementary Materials*).

## Ethics Statement

The research protocol was reviewed and approved by the Ethics Committee of Anhui Institute of Medicine (Approval No. 2023-LLBG-024). All animal experiments were carried out in compliance with institutional guidelines and received approval from the Experimental Animal Ethics Committee of Anhui Academy of Medical Sciences (Approval No. AHAMS-AER-AHGXKY-2025-0007).

## Data Availability Statement

All relevant data are within the manuscript and its Supporting Information files.

## Acknowledgments

The authors sincerely thank the Department of Pathology at the Affiliated Hospital of Anhui Institute of Medicine (Hefei, China) for providing HCC tissue specimens and the corresponding anonymized clinical records used in this study.

## Author Contributions

Y.Z.: conceptualization, methodology, designed the experiments, analysis. X.W.: validation, pathological assessment. X.P.: samples collection and data collection. Y.Z. and X.P.: funding support. All authors agreed the manuscript for publication.

